# Identification of ER:melanosome membrane contact sites in the retinal pigment epithelium

**DOI:** 10.1101/2025.01.07.631750

**Authors:** T. Burgoyne, D. Doncheva, E.R. Eden

## Abstract

The retinal pigment epithelium (RPE) forms a monolayer of cells at the blood:retina interface that plays important roles for photoreceptor renewal and function and is central to retinal health. RPE pigment is provided by melanin-containing melanosomes which offer protection against light and oxidative stress. Melanosome migration into the apical processes of the RPE following light onset is thought to contribute to preventing retinal degeneration with age, though the mechanism is not yet clear. Melanosomes are transported along microtubules to the apical surface where they are transferred to actin filaments within the apical processes. Melanosomes are lysosome-related organelles derived from endosomes and endosome transport along microtubules is heavily influenced by the endoplasmic reticulum (ER) through ER:endosome contact sites. Here we describe extensive connection between the ER and melanosomes in the RPE. We further show, in skin melanocytes, that the ER forms contact sites with all stages of melanosome maturation, but ER contact is reduced as melanosomes mature. Finally, we identify tripartite contact sites between the ER, melanosomes and mitochondria in both RPE tissue and cellular models, suggesting that the ER may influence melanosome biogenesis, maturation and interaction with mitochondria.

## Introduction

The pigment melanin provides photoprotection to the skin and eyes by absorbing and scattering ultraviolet radiation and protecting against oxidative stress. Melanin is produced and stored within melanosomes, membrane-bound organelles found in skin epidermal melanocytes and in the retinal pigment epithelium (RPE) cells of the eye. Described as lysosome-related organelles, melanosomes originate from endosomes but undergo a series of maturation stages (I to IV), involving the progressive deposition of melanin pigment^1^. Defects in melanosome biogenesis/function is associated with several human diseases including melanomas and often causes albinism and loss of vision^2^.

The RPE forms a monolayer of cells at the blood-retina interface at the back of the eye. Melanogenesis occurs prenatally in the RPE, which is densely packed with mature melanosomes from birth, providing cells with their characteristic pigmentation and offering protection against harmful backscattered light and damaging reactive oxygen species (ROS)^3^. Melanosome distribution in the RPE is regulated, at least in part, by the light cycle. Light onset triggers apical migration of melanosomes along microtubules and their transfer to the actin filaments of the apical processes that extend between photoreceptor outer segments^4^.

In contrast to the RPE, in skin and hair, melanosome biogenesis occurs continuously in melanocyte cells, predominantly in a central perinuclear region of melanocytes. Mature melanocytes traffic towards the cell periphery and along dendrites that extend through the epidermis to keratinocytes. Subsequent exocytosis from melanocyte dendrites releases the melanosome core for phagocytosis into keratinocytes^5^. Melanin-containing phagosomes ultimately fuse with the lysosome to form “melanokerasomes”, weakly acidic terminal organelles that cluster around the nucleus forming melanokerasome:nuclear contact sites to provide a photoprotective perinuclear “cap” that protects DNA from UV damage^6^.

Although melanocyte distribution and its regulation differ between melanosomes and the RPE, basic mechanisms that drive melanosome movement are common to both cell types. Melanosomes are transported along microtubules to the periphery of the cell where they interact with the actin cytoskeleton via tripartite protein complexes consisting of Rab27A, it’s effector and an actin motor protein (Myosin-7a in the RPE and Myosin-V in melanocytes). In non-polarised cells, including melanocytes, the microtubule-organizing centre (MTOC), in which the minus ends of microtubules are anchored, is in a perinuclear position. In polarised RPE however, the MTOC is apically positioned, with microtubules extending vertically between minus ends in the apical region and plus ends facing the basal surface^7^. Transport along microtubules can be bidirectional, with anterograde transport towards plus ends (ie, at the periphery in melanocytes/basal surface in RPE) mediated by kinesin motor proteins, while retrograde transport towards the minus end (at the cell centre in melanocytes or apical side in RPE) is mediated by dynein motor proteins^8^. Thus dynein-dependent transport delivers mature melanosomes to the actin-rich apical domain of RPE cells but favours perinuclear positioning in melanocytes^9^.

Similarly, distribution of endosomes, from which melanosomes are derived, is also dependent on dynein and kinesin but endosome interactions with microtubule motor proteins is strongly influenced by the ER through ER:endosome membrane contact sites (MCS). In complementary, and likely coordinated processes, MCS in non-polarised cells promote kinesin-loading for plus-end directed endosome movement along microtubules to the plasma membrane^10^ and prevent dynein interaction with endosomal RILP to reduce minus-end directed movement towards the peri-nuclear compartment^11^. ER:endosome MCSs thereby couple the regulation of both plus and minus-end-directed transport and interestingly ER contact with melanosomes has been identified in pigmented keratinocytes^12^.

Melanosomes have also been shown to form contact sites with another organelle in melanocytes, where physical bridging with mitochondria has been implicated in melanosome biogenesis. These mitochondria:melanosome MCS are tethered by Mitofusin-2, are enhanced by the ocular albinism type 1 (OA1) protein that can stimulate melanogenesis and are increased in the perinuclear region where melanosomes are generated^13^. Melanosome biogenesis is heavily influenced by mitochondria, which likely supply ATP to meet the energy needs of melanogenesis, as well as buffering cytosolic Ca^2+^ since both the mitochondrial calcium uniporter (MCU) that mediates Ca^2+^ influx into mitochondria^14^ and the mitochondrial voltage-dependent anion channel, VDAC1^15^ were recently found to be important in the regulation of melanogenesis.

Here we identify MCS between melanosomes and the ER both in RPE tissue and cellular models, from multiple species. Characterisation of ER:melanosome MCS reveals that the proportion of melanosomes forming an ER contact site is conserved across species and that the ER contacts all stages of melanosome maturation in melanocytes. Our data further suggests the existence of tripartite connections between the ER, melanosomes and mitochondria.

## Results

### Melanosomes contact the ER and mitochondria in mouse, porcine and human RPE

To determine if melanosomes in the RPE form ER contact sites, we imaged mouse retina by electron microscopy (EM). Electron-dense pigmented melanosomes were readily visible both in central areas and apical processes of the RPE from mouse eyes two hours after light onset (Figure 1A), consistent with the previously described Rab27a-dependent increased apical positioning of melanosomes in response to light^4^. Just as melanosome:mitochondria MCS have been identified in melanocytes^13^, we also found mitochondria tightly associated with melanosomes in mouse RPE (Figure 1A; mitochondria at MCS false-coloured green). Both the subpopulation of mitochondria in contact with the basal infoldings of the RPE^16^ (Figure 1A, arrows), and mitochondria in central/apical regions (Figure 1A) appear to form contact sites with melanosomes.

**Figure 1:**
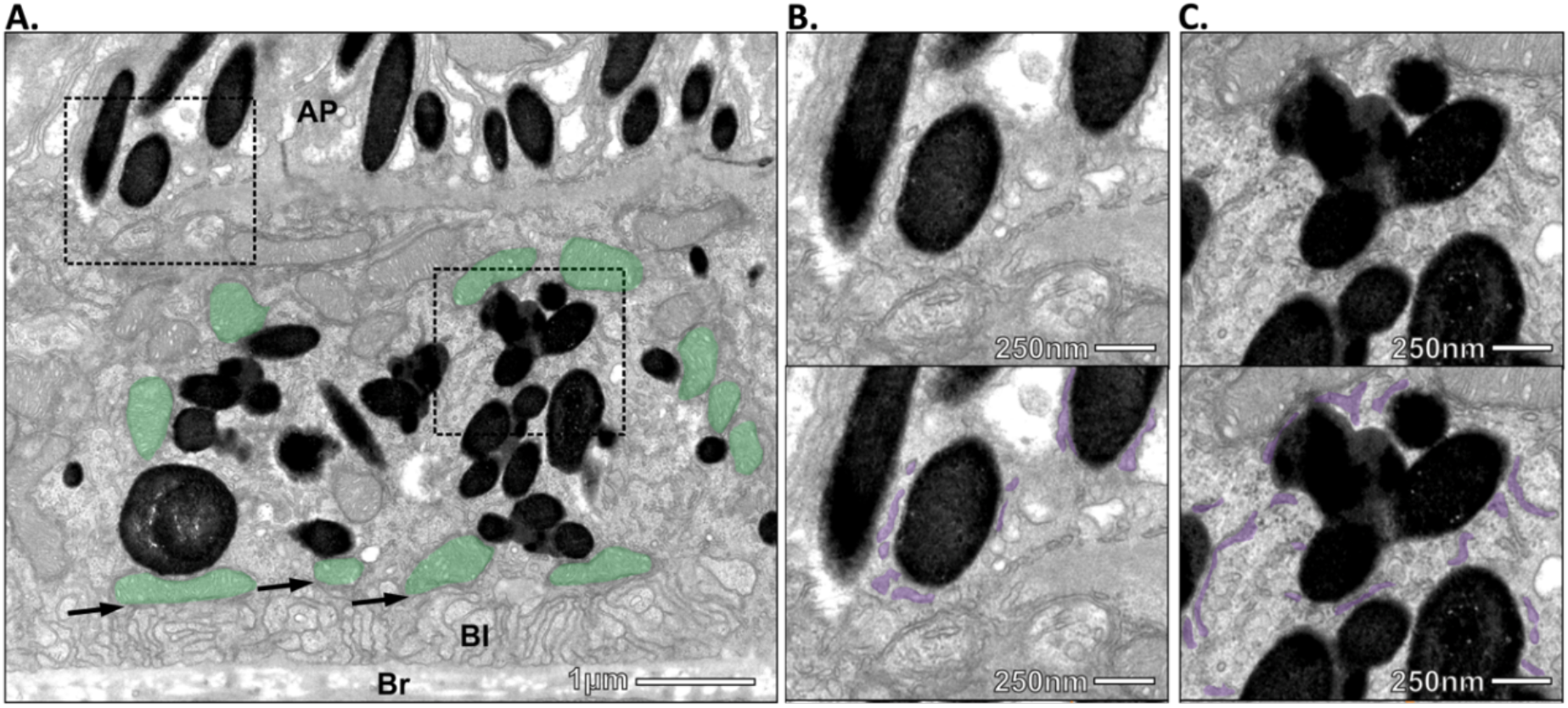
ER:melanosome MCS in RPE from adult mouse tissue. Ultrathin sections through the retina of mouse eyes were imaged by transmission electron microscopy. **A)** Highly pigmented, electron dense melanosomes form extensive MCS with ER, both in the apical (upper box) and central (lower box) areas of the RPE and with mitochondria in central and basal areas (mitochondria at MCS false-coloured green). Arrows, mitochondria contacts with basal infoldings. Br, Bruch’s membrane; Bl, basal infoldings; AP, apical processes. Scale bar, 1μm Higher magnification images of apical (B) and central C) areas are shown with the ER false-coloured lilac in the lower panel. Scale bar, 250nm.

Higher magnification imaging revealed that in addition to mitochondrial contact, melanosomes also form abundant connections with the ER, in both apical (Figure 1B) and central (Figure 1C) regions of mouse RPE. To determine if this RPE contact site population is unique to mouse tissue, we next examined RPE from human and porcine tissue. ER:melanosome MCS were readily visible in RPE from both species (supplemental Figure S1) indicating that formation of ER:melanosome MCS is conserved across different species and likely play important functional roles.

Having identified ER:melanosome MCS in mouse, porcine and human RPE tissue, we next investigated their formation in cellular RPE models in culture. Primary porcine RPE have been widely used to successfully generate pigmented polarized epithelia^17^ and we found abundant ER:melanosome MCS visible by EM in primary porcine RPE cells (supplemental Figure S2A), comparable to the extent found in *in vivo* models (51% melanosomes had an ER contact in primary porcine RPE compared with 54% in RPE from mouse tissue).

**Figure S1:**
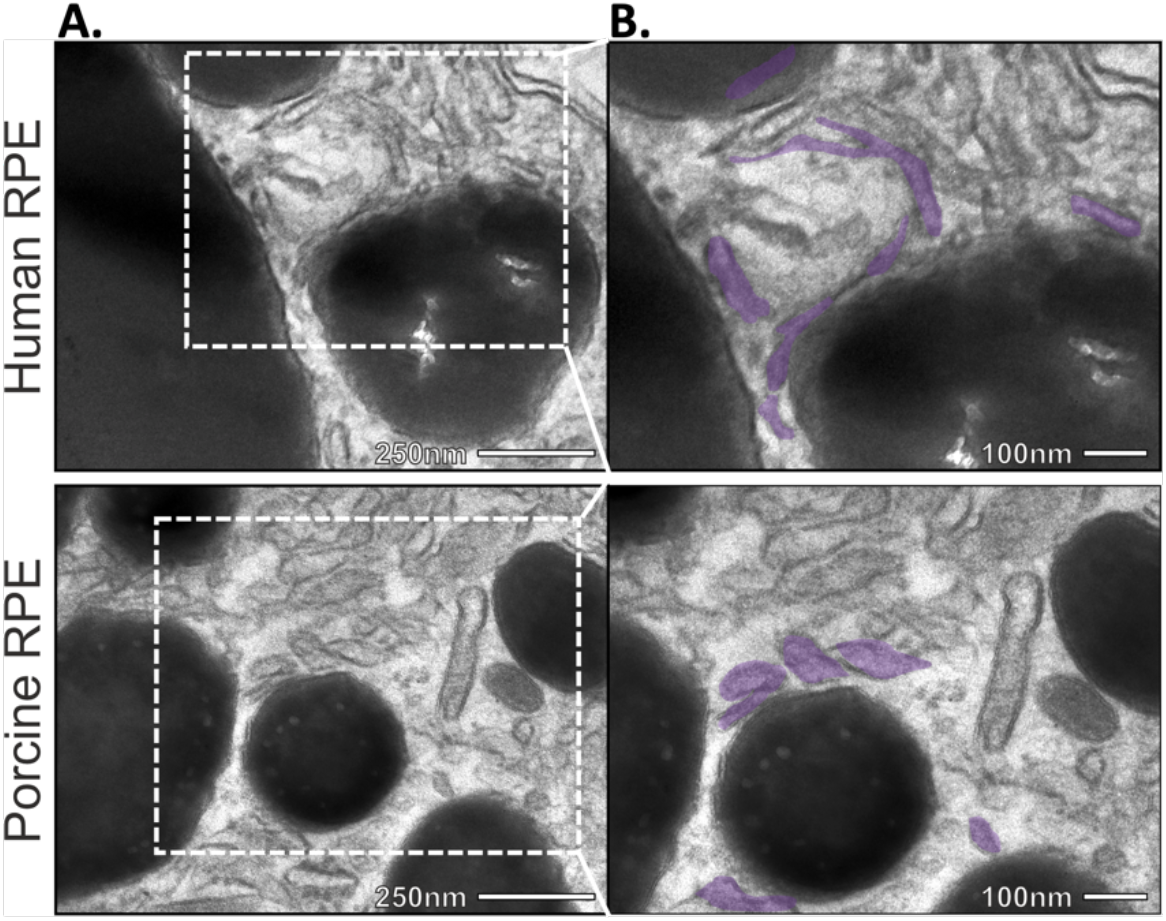
ER:melanosome MCS in RPE from human and porcine tissue. **A**) Ultrathin sections through the retina of human (upper panel) or porcine (lower panel) eyes were imaged by transmission electron microscopy. Scale bar, 250nm. **B**) Higher magnification of boxed regions within A, with the ER at MCS false-coloured lilac. Scale bar, 100nm.

The human ARPE19 cell line is a widely used RPE model that can be differentiated into RPE-like cells^18^. Following six weeks of differentiation in X-Vivo medium, wild-type ARPE-19 cells formed a highly pigmented monolayer and analysis by EM revealed that ER:melanosome MCS also form in differentiated ARPE-19 cells (supplemental Figure S2B) to an even greater extent, with 74% melanosomes forming an ER contact.

### ER:melanosome MCS form during foetal development in mouse RPE

Biogenesis and maturation of melanosomes in the RPE occurs during foetal development in mice^19^. We therefore examined melanosome MCS in pre-existing sections from E14 foetal mouse eyes (not yet exposed to light). Melanosome synthesis is thought to be completed before birth in mouse RPE, with pigment granules retained throughout life^20^ and our data indicate that melanosome biogenesis and maturation has occurred by E14 during foetal development in mice (Figure 2). Using electron tomography, we were able to identify discreet ER contact sites with mature melanosomes in E14 mouse RPE, even though there appeared to be less ER content compared with postnatal RPE (Figure 2A). The ER leading to the melanosome contact was often decorated with ribosomes (false-coloured blue in Figure 2A), indicative of rough ER, though the ribosomes appeared to be excluded from the MCS, as has been reported at other ER contact sites^21^. In some areas, thin electron-dense strands tethering the two organelles could be discerned (Figure 2B), suggestive of a functional population of MCS that are stabilised by tethering complexes.

**Figure S2:**
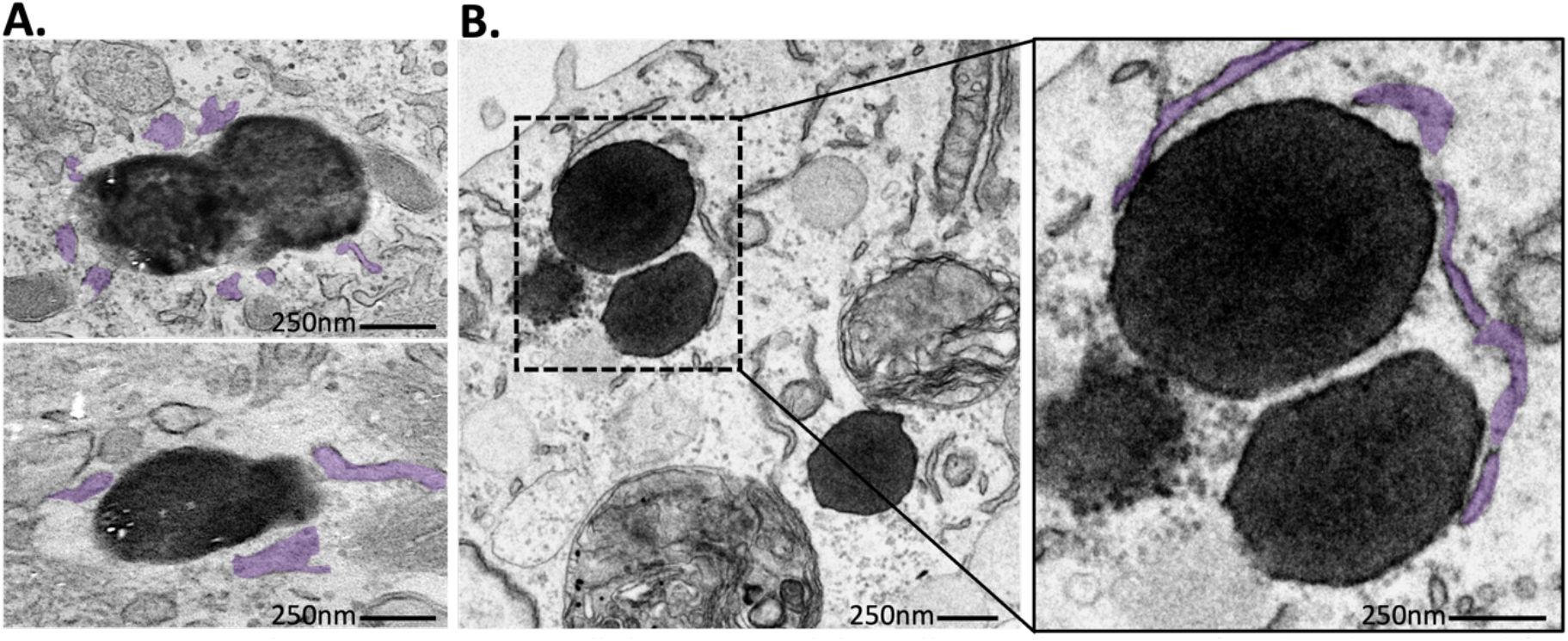
ER:melanosome MCS in cellular RPE models. Cells in culture were fixed and prepared for electron microscopy. **A**) Primary porcine RPE isolated from porcine eye tissue and passaged once onto transwell membranes prior to fixation. Images show ER at MCS false-coloured lilac. **B**) ARPE19 cells were cultured in X-Vivo medium for 6 weeks to gain pigmentation. Magnified images of the boxed region is shown on the right with the ER at MCS false-coloured lilac. Scale bar, 250 nm.

**Figure 2:**
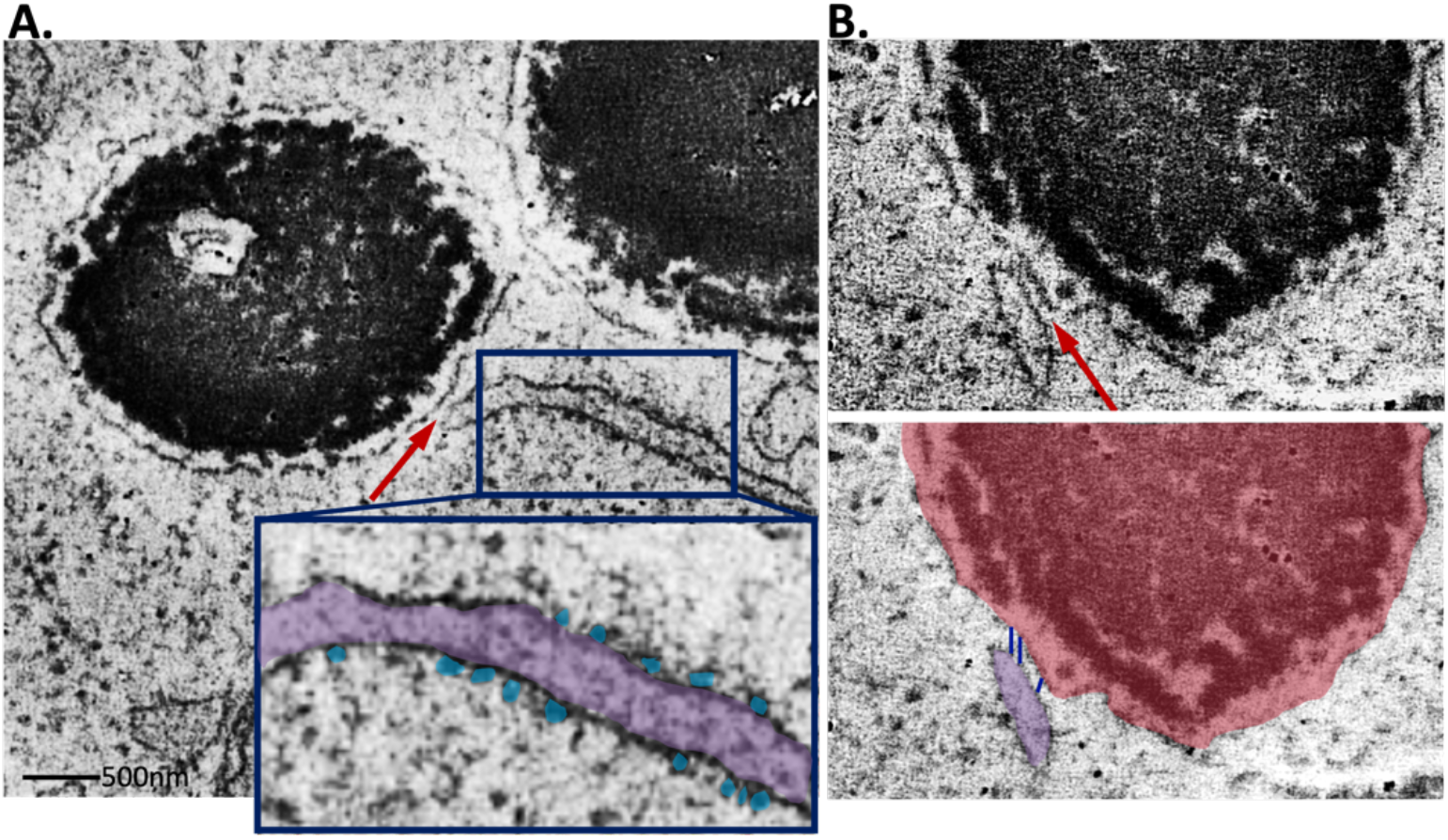
Electron tomograph of RPE from E14 mouse tissue. The melanosome limiting membrane surrounding the melanocore forms ER contact sites (red arrows). **A**) The ER (false-coloured lilac in the magnified boxed region) is decorated with ribosomes (false-coloured blue) which appear excluded from the site of contact with the melanosome limiting membrane (red arrow). Scale bar, 500 nm. **B**) Inter-organelle tethers (false-coloured navy in the lower panel) at the contact site between ER (lilac) and melanosomes (burgundy).

### ER:melanosome MCS form with all stages of melanosome maturation in human melanocytes

Whereas melanosome biogenesis and maturation is largely complete before birth in RPE cells, in skin melanocytes, melanosome synthesis occurs continuously throughout life^20^. Thus, melanocytes provide the opportunity to examine all four maturation stages and to determine the extent of ER contact with early as well as mature melanosomes. We therefore imaged different stages of melanosome maturation in the human melanoma MNT-1 cell line. In addition to forming abundant MCS with mature (stage IV) melanosomes as seen in RPE cells, the ER also made connections with melanosomes at earlier stages (I-III) of maturation (Figure 3A). In contrast to the correlation between endosome maturation and extent of ER contact^22^, we found increased ER contact with early stage melanosomes compared with more mature organelles (Figure 3B), suggesting an inverse correlation between melanosome maturation and extent of ER contact.

**Figure 3:**
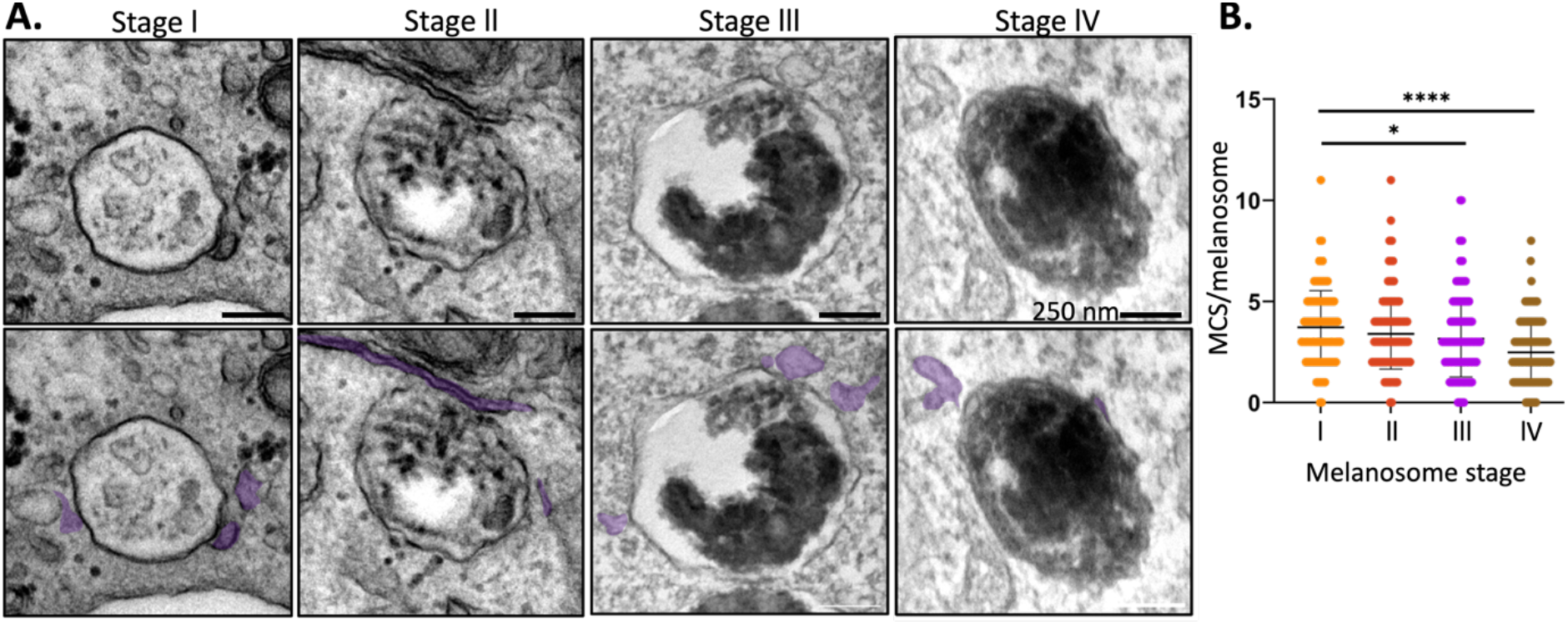
The ER forms MCS with all melanosome maturation stages. MNT-1 melanocytes were fixed and prepared for EM. **A**) Electron micrographs show ER (false-coloured lilac on the lower panels) interacting with melanosomes at all four stages of maturation. Scale bar, 250 nm. **B**) Quantitation of the average number of ER contacts/melanosome at each stage of maturation from 2 independent experiments with statistical significance determined by one-way ANOVA, Brown-Forsythe and Welch multiple comparison.

### Tripartite MCS between ER, melanosomes and mitochondria

In some areas of mitochondria:melanosome association, the ER appears to run intermittently between the two organelles, while in other areas, ER:melanosome contact is directly adjacent to mitochondria:melanosome MCS (Figure 4), suggesting the existence of tripartite contact sites between the three organelles. These three-way ER:melanosome:mitochondria contacts were present in both mouse tissue RPE (Figure 4A) and primary porcine RPE cells (Figure 4B), but were less frequent in mouse RPE, where 7% melanosomes were in a tripartite MCS compared with 13% in primary porcine RPE. This could reflect structural differences between *in vitro* and *in vivo* RPE models since a significant proportion of melanosomes migrate into the apical processes that interdigitate between photoreceptor outer segments following light onset, *in vivo*. In contrast, apical processes are much less pronounced *in vitro* and do not contain melanosomes. Distribution of mitochondria, like melanosomes, is also influenced by light. The majority of mitochondria localise to baso-lateral RPE, where a proportion are anchored to RPE basal infoldings through mitochondria:plasma membrane MCS^16^, while light onset stimulates migration of a different mitochondrial subset into central RPE towards the apical surface, but not into apical processes^16^. The mouse eyes were fixed two hours after light onset but it is likely that the number of tripartite MCS in mouse RPE might be higher without light-stimulated transport of melanosomes into the apical processes. Together, these data suggest that approximately 10% of melanosomes form multi-organelle platforms with the ER and mitochondria for non-vesicular communication and exchange.

**Figure 4:**
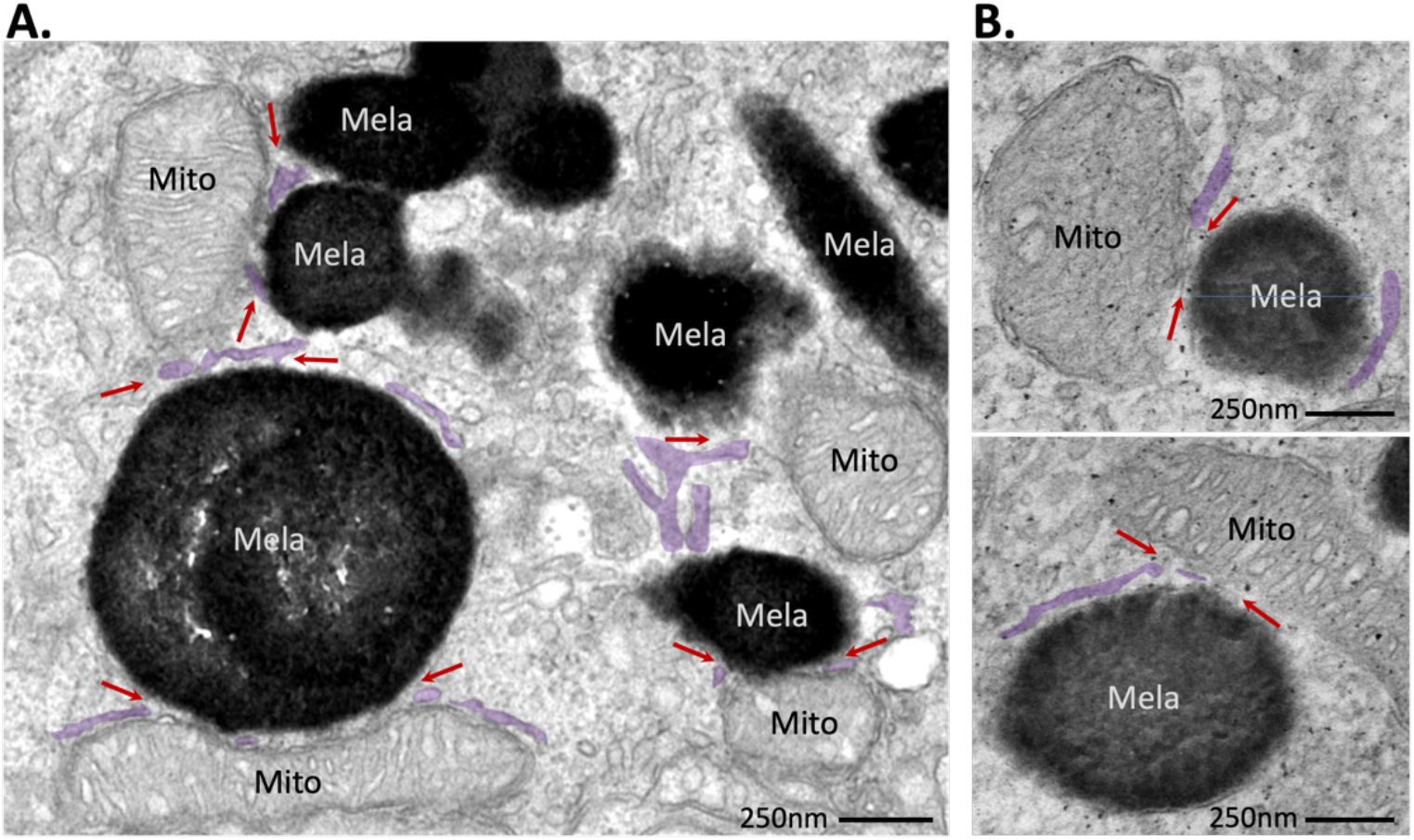
Tripartite MCS between ER, melanosomes and mitochondria. Electron micrographs of RPE showing ER:melanosome:mitochondria MCS (arrows). ER, false-coloured lilac; melanosomes, Mela; mitochondria, Mito; scale bar, 250 nm. **A**) RPE from adult mouse tissue. **B)** Primary porcine RPE cells.

## Discussion

We have identified extensive ER contact with melanosomes that are conserved across species and in both RPE tissue and cells in culture. The frequency and extent of these MCS suggest important physiological roles which are as yet unknown. Melanosome traffic into the apical processes of the RPE is important in preventing retinal degeneration^23^, but although ER contact has been shown to regulate the positioning of endosomes^24^, from which melanosomes are derived, the influence of the ER over melanosome distribution has not yet been explored. Whereas in non-polarised melanocyte cells the microtubule-organizing centre (MTOC) is typically peri-nuclear, the MTOC is in the apical region of polarised RPE cells. Thus the minus-end directed motor protein dynein moves melanosomes away from dendrites, towards the cell body of melanocytes, while in the RPE, dynein-mediated transport traffics melanosomes in the opposite direction, to the apical surface where they are retained by myosin-7a/RAB27A/MYRIP-dependent transfer to actin filaments^9^. Endosomal engagement of dynein for minus-end directed transport along microtubules is negatively regulated by ER contact. The endosomal sterol-binding protein ORP1L forms a complex with the small GTPase Rab7 and the Rab7 effector RILP. In the absence of ER contact, the p150Glued subunit of the dynein/dynactin microtubule motor complex directly binds RILP, mediating minus-end-directed transport. However, interaction of ER-localised VAP with ORP1L at ER:endosome MCS results in removal of p150Glued and dynein, preventing dynein-dependent minus-end directed transport^11^. Our data has revealed that the majority (over 50%) of mature melanosomes form an ER contact in mouse RPE two hours after light onset, when maximal melanosome positioning in apical processes is expected^4^. This suggests a potential role for ER contact sites in returning melanosomes from apical processes to the cell body and in restricting further dynein-dependent transport to the apical domain of the RPE. If MCS orchestrate melanosome positioning, the light-dependent migration of melanosomes into apical processes would suggest that light exposure may have some influence over contact site formation.

Whereas melanosome biogenesis and maturation are mostly prenatal in the RPE, in melanocytes it is continuous throughout life. Interestingly ER contact with early melanosomes was more extensive than with more mature organelles in melanocytes, suggesting a possible role for the ER in melanosome biogenesis/maturation, likely through lipid and/or Ca^2+^ flux as has been described at ER:endosome MCS^24^. After six weeks of differentiation in X-Vivo medium, ARPE19 cells continued to gain pigment, suggesting that melanosome maturation was incomplete, which may explain the greater extent of ER contact in these cells (74% melanosomes formed an ER contact compared with 54% in mouse RPE) since we found that ER contact is increased with less mature organelles.

The tripartite ER:melanosome:mitochondria MCS may also point to a role for the ER in melanosome biogenesis/maturation as has been shown for mitochondria. Melanosome biogenesis is subject to regulation by microphthalmia-associated transcription factor (MITF), a key transcription factor for expression of melanogenic enzymes^25^ and increased cytosolic Ca^2+^ on loss of mitochondrial VDAC1 induced downstream MITF activation and melanosome biogenesis^15^. Lysosome MCS with ER and mitochondria have both been implicated in contributing to Ca^2+^ flux^26-29^ and melanosome Ca^2+^ import from mitochondria has been implicated in melanosome biogenesis^30^. Thus, these tripartite contacts may act as Ca^2+^ signalling hubs to regulate melanosome biogenesis. An additional role for melanosomes in quenching mitochondrial ROS has also been proposed^13, 31^; the ER may help to coordinate melanosome:mitochondria contact through tripartite MCS to facilitate ROS scavenging by melanosomes. Identifying the molecular composition of melanosome MCS tethering complexes will be key to elucidating their function in the RPE.

## Methods

### RPE tissue

Mouse tissue was isolated as previously described^16^, with mice sacrificed 2h after light onset and eyes immediately plunged into EM fixative for 1h prior to dissection. Human eye tissue from a donor with no known eye disease was approved for research and received from the Moran Eye Institute, Salt Lake City, UT, USA, courtesy of Dr GS Hageman. Porcine eyes were collected in the morning after light onset, kept on ice during transport and fixed for electron microscopy in the afternoon.

### Cell culture

Primary porcine RPE cells were isolated from porcine eyes as previously described^32^ and seeded into 6 well plates. Once fully confluent, cells were passaged onto Corning transwell cell culture inserts. Cells were cultured in DMEM Glutamax supplemented with 1% FBS/PS. ARPE19 cells were passaged onto transwells and cultured in X-Vivo medium (Lonza Bioscience) for 6 weeks until they formed a pigmented cobblestone monolayer.

### Electron microscopy

The cornea and lens were removed from all eyes prior to subsequent incubation in EM fixative (2% gluteraldehyde, 2% paraformaldehyde in 0.1M sodium cacodylate buffer) for 1 hour (mouse tissue), overnight (porcine tissue) or >24h, until further processing (human tissue). Cells in culture were fixed in EM fixative for 30 min. Following fixation, all samples were prepared for EM as previously described^16^.

